# ChromaFold predicts the 3D contact map from single-cell chromatin accessibility

**DOI:** 10.1101/2023.07.27.550836

**Authors:** Vianne R. Gao, Rui Yang, Arnav Das, Renhe Luo, Hanzhi Luo, Dylan R. McNally, Ioannis Karagiannidis, Martin A. Rivas, Zhong-Min Wang, Darko Barisic, Alireza Karbalayghareh, Wilfred Wong, Yingqian A. Zhan, Christopher R. Chin, William Noble, Jeff A. Bilmes, Effie Apostolou, Michael G. Kharas, Wendy Béguelin, Aaron D. Viny, Danwei Huangfu, Alexander Y. Rudensky, Ari M. Melnick, Christina S. Leslie

## Abstract

The identification of cell-type-specific 3D chromatin interactions between regulatory elements can help to decipher gene regulation and to interpret the function of disease-associated non-coding variants. However, current chromosome conformation capture (3C) technologies are unable to resolve interactions at this resolution when only small numbers of cells are available as input. We therefore present ChromaFold, a deep learning model that predicts 3D contact maps and regulatory interactions from single-cell ATAC sequencing (scATAC-seq) data alone. ChromaFold uses pseudobulk chromatin accessibility, co-accessibility profiles across metacells, and predicted CTCF motif tracks as input features and employs a lightweight architecture to enable training on standard GPUs. Once trained on paired scATAC-seq and Hi-C data in human cell lines and tissues, ChromaFold can accurately predict both the 3D contact map and peak-level interactions across diverse human and mouse test cell types. In benchmarking against a recent deep learning method that uses bulk ATAC-seq, DNA sequence, and CTCF ChIP-seq to make cell-type-specific predictions, ChromaFold yields superior prediction performance when including CTCF ChIP-seq data as an input and comparable performance without. Finally, fine-tuning ChromaFold on paired scATAC-seq and Hi-C in a complex tissue enables deconvolution of chromatin interactions across cell subpopulations. ChromaFold thus achieves state-of-the-art prediction of 3D contact maps and regulatory interactions using scATAC-seq alone as input data, enabling accurate inference of cell-type-specific interactions in settings where 3C-based assays are infeasible.

---

Genome-wide chromosome conformation capture techniques such as Hi-C, HiChIP, and ChIA-PET^1–3^ provide powerful tools for mapping cell-type-specific regulatory interactions that can link enhancers to genes and enable the interpretation of non-coding disease-associated variants^4, 5^—at least when there is sufficient input material to generate high-complexity libraries and allow for very deep sequencing. Indeed, the use of these assays is often impeded by their substantial costs, time requirements, and technical difficulty, especially when studying rare cell populations where obtaining a sufficient number of cells for a high-quality contact map becomes impractical^6, 7^. On the other hand, single-cell chromosome conformation mapping technologies, such as single-cell Hi-C or ChIA-Drop, although exciting, are experimentally challenging and produce sparse data sets that are typically analyzed at 100kb-1Mb resolution^8–11^. By contrast, single-cell chromatin accessibility (scATAC-seq) datasets can be readily generated from small amounts of input material due to the availability of commercial kits^12^. Genome-wide chromatin accessibility profiles reflect the extent to which nuclear molecules, including transcription factors, chromatin remodelers, histones and other chromatin-associated proteins, can physically interact with chromatinized DNA, and single-cell chromatin accessibility contains subtle information about pairs of interacting loci that are jointly associated with DNA-bound factors^13^. This raises the question of whether one can predict chromatin interactions and connect regulatory elements to their target genes using scATAC-seq data alone.

Several models have been proposed to predict chromatin interactions from genomic sequence and easier-to-obtain bulk or single-cell epigenomic data^14–19^. For instance, Cicero was the first method to leverage the co-accessibility structure between accessible elements (‘peaks’) in scATAC-seq data to infer chromatin interactions in an unsupervised fashion^18^. DeepC^19^, Akita^14^ and Orca^15^ are supervised deep neural network-based models that predict chromatin contact maps from genomic DNA sequence. Epiphany, a model we introduced recently for cell-type-specific contact map prediction, uses a collection of bulk 1D epigenomic input tracks to enable generalization to novel cell types^17^. Another recent model, C.Origami, is also capable of making cell-type-specific predictions using DNA sequence together with bulk ATAC-seq and CTCF ChIP-seq in the target cell type^16^. However, these existing models for chromatin interaction prediction have practical limitations. Unsupervised models like Cicero offer modest accuracy, whereas sequence-based models such as DeepC, Akita and Orca fail to generalize effectively to new cell types and indeed tend to predict similar contact maps across training cell types^14, 16^. Meanwhile, C.Origami and Epiphany both require multiple input data modalities, which are not always available, and C.Origami in particular employs a more complex model that may be susceptible to overfitting^20^.

In this study, we introduce ChromaFold, a supervised deep learning model that predicts the 3D contact map from scATAC-seq data and CTCF motif tracks as input features. Given the linkage between the accessibility landscape of regulatory elements and 3D genome organization, our underlying hypothesis is that we can leverage the covariation in accessibility stemming from asynchronous chromatin looping events across single cells. This assumption is further substantiated by prior studies showing that pairs of genomic bins with high co-accessibility are enriched for chromatin looping events^18, 21^. Additionally, given the crucial role of the CTCF protein in shaping 3D chromatin structure, inclusion of CTCF-associated signals is expected to enhance the model’s predictive power^22–24^. For wider adaptability, we do not require CTCF ChIP-seq as an input and offer two versions of ChromaFold. *ChromaFold +CTCF motif* uses CTCF motif score, a measure of the likelihood that a genomic region contains a binding site for the CTCF protein, as a proxy for CTCF binding^25^. *ChromaFold +CTCF ChIP* uses the actual CTCF ChIP-seq track as input (unless otherwise noted, ChromaFold refers to *ChromaFold +CTCF motif*).

The key advantages of ChromaFold include its requirement of only scATAC-seq data as experimental input data, its ability to make cell-type-specific predictions in new cell types, and its lightweight architecture, making it compatible with standard GPUs. Importantly, ChromaFold can also be employed to deconvolve bulk chromatin interaction data across constituent cell types— resolving the cell-type-specificity of chromatin interactions—by fine-tuning on bulk Hi-C and scATAC-seq data from the same complex tissue.

We evaluated ChromaFold on five human and three mouse test cell types and tissues. ChromaFold was able to make accurate cell-type-specific predictions of 3D contact maps (as evaluated by distance stratified Pearson correlation) and peak-level interactions (as evaluated by receiver operating characteristic and precision-recall analysis) in new cell types and species. In particular, ChromaFold predictions at important lineage-defining loci in murine germinal center B cells (GCBs), regulatory T (Treg) cells, and hematopoietic stem cells (HSCs) recovered correct cell-type-specific regulatory interactions. Interestingly, despite its smaller model and reduced information requirements, ChromaFold’s performance was comparable to C.Origami when using predicted CTCF motif information as input and outperformed C.Origami when using CTCF ChIP-seq track as input on new cell types. Finally, using paired Hi-C and scATAC-seq in human pancreatic islets, ChromaFold successfully deconvolved chromatin interactions into those specific to alpha cells and beta cells.

Overall, ChromaFold achieves state-of-the-art generalization to novel cell types while requiring only a single input modality to enable accurate contact map and regulatory interaction prediction in any setting where scATAC-seq can be generated.

## Results

### ChromaFold is a deep learning model that predicts 3D contact maps from scATAC-seq data

To enable fast and accurate prediction of chromatin contacts from scATAC-seq data alone, we developed ChromaFold, a lightweight convolutional neural network-based model that makes cell-type-specific predictions. ChromaFold is trained on paired scATAC-seq and Hi-C data from a panel of training cell types. ChromaFold takes three input tracks—pseudobulk chromatin accessibility and correlation structures in accessibility (co-accessibility) profiles across cells, both computed from scATAC-seq, and predicted CTCF motif scores—all processed for a 4.01Mb genomic region (**Fig. 1a**). These processed inputs are passed through two feature extractors in the ChromaFold architecture. The first feature extractor takes the pseudobulk accessibility and CTCF motif score tracks as input, while the second takes the co-accessibility as input. For memory efficiency, we only compute the co-accessibility between the genomic bins in the center 10kb region with the rest of the bins in the 4.01Mb region as input. These extractors produce a latent representation of the genomic region, which is then passed through the linear predictor to predict the chromatin interactions between the center genomic bin and its neighboring bins within a 2Mb distance (V-stripe) at 10kb resolution, using the HiC-DC+ Z-score^26^ normalized Hi-C contact map for the corresponding region and cell type as the target (**Fig. 1b**, **Supplementary Fig. 1a**).

**Figure 1.**
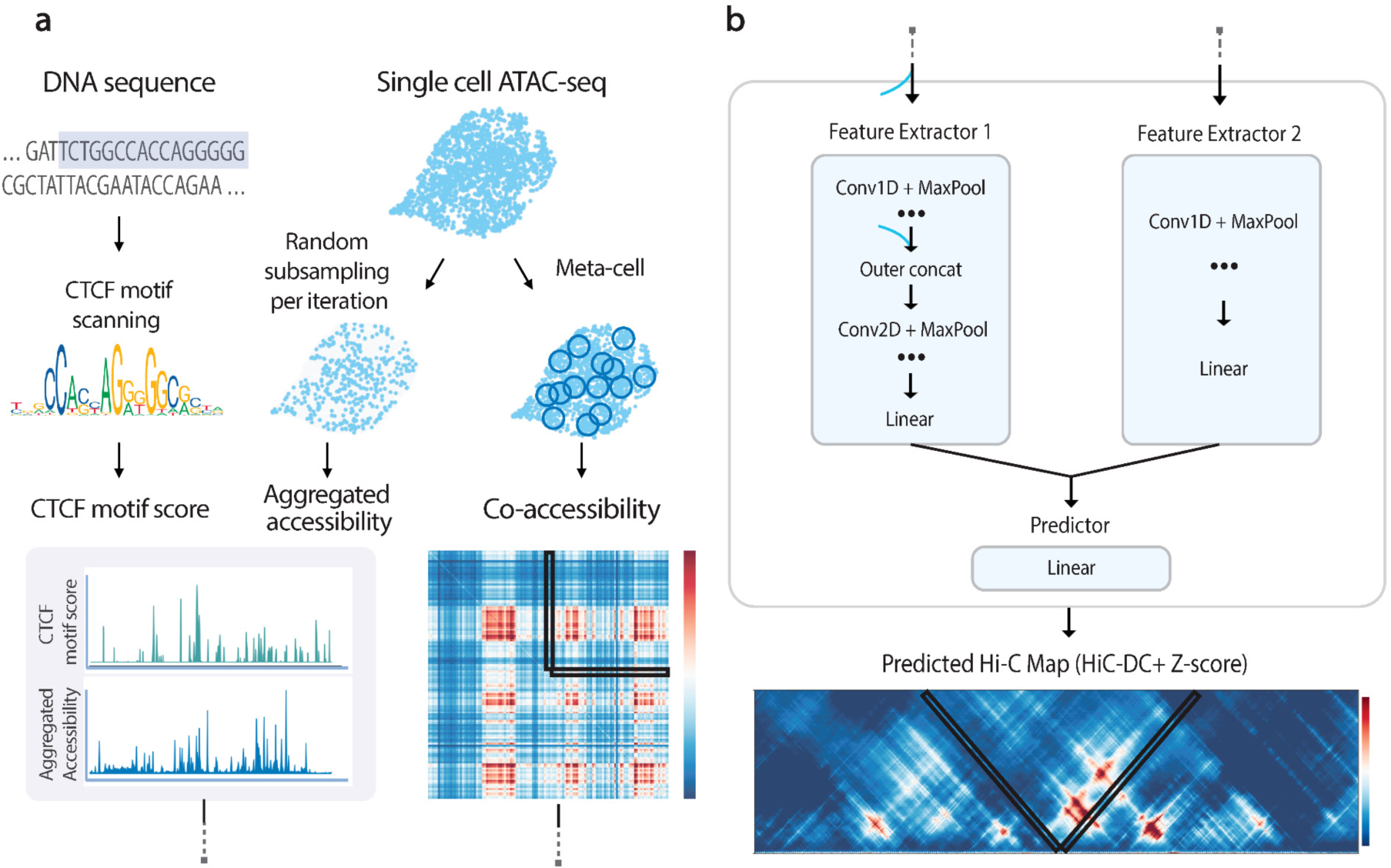
ChromaFold predicts the 3D contact map from scATAC-seq alone. ChromaFold is a deep learning model that enables prediction of 3D contact maps solely from scATAC-seq data, using pseudobulk chromatin accessibility and co-accessibility from scATAC-seq as well as predicted CTCF motif tracks as input features. **a.** Schematic of the ChromaFold input data processing framework. **b.** ChromaFold model architecture. The model consists of two feature extractors: feature extractor 1 for the aggregated accessibility and CTCF motif score tracks, and feature extractor 2 for the co-accessibility extracted from a V-stripe region. The feature extractors produce a latent representation of the 4Mb genomic region. The Z-score predictor then takes this latent representation and predicts the chromatin interactions between the center genomic tile and its neighboring bins within a 2Mb distance, annotated by the V-shaped black box. Each genomic tile is 10Kb in length.

To process the input data, the CTCF motif score track is generated by scanning a set of CTCF position weight matrices^27, 28^ (**Supplementary Fig. 1b**) across the DNA sequence. The pseudobulk chromatin accessibility is obtained by aggregating the accessibility profile across single cells in a population. The co-accessibility is derived by first generating metacells to combat sparsity, then calculating the Jaccard similarity^29^ between binarized accessibility profiles across metacells. During training, we randomly subsample single cells and metacells from the population per iteration to generate pseudobulk accessibility and co-accessibility input data, respectively. This data augmentation step is critical for improving model generalizability to data sets of varying quality and size^30–32^. As a sanity check, we observed an enrichment of CTCF occupancy as measured by ChIP-seq in genomic bins with high CTCF motif score (**Supplementary Fig. 1c**), and an enrichment of chromatin interactions as measured by Hi-C in co-accessible genomic bins for datasets with greater variability (**Supplementary Fig. 1d**). These results suggest that our input tracks provide valuable information for predicting chromatin contacts that can be harnessed by ChromaFold when trained across sufficiently diverse training cell types.

We trained ChromaFold on three human cell types (IMR-90, GM12878, and HUVEC) to improve model generalizability to novel test cell types. Fifteen chromosomes were used for training, two for validation, and four held-out for testing and evaluating model performance. We held out three other human cell types (K562, hESC and activated CD4+ T cells) to test how well ChromaFold can generalize to new cell types. The full contact map was obtained by combining the V-stripe predictions along the chromosome (**Methods**). To evaluate ChromaFold’s performance, we assessed both the chromosome-wide contact map and significant interaction prediction (based on HiC-DC+ top-scoring interactions) on held-out chromosomes for both training and held-out cell types (**Fig. 2a,b**). ChromaFold achieved an average distance-stratified Pearson correlation of 0.55-0.60 and 0.45-0.47 and average area under the ROC curve (AUROC) of 0.84-0.85 and 0.77-0.79 in training and held-out cell types, respectively. These results demonstrate ChromaFold’s ability to effectively predict the 3D contact map in unseen data and capture significant interactions.

**Figure 2.**
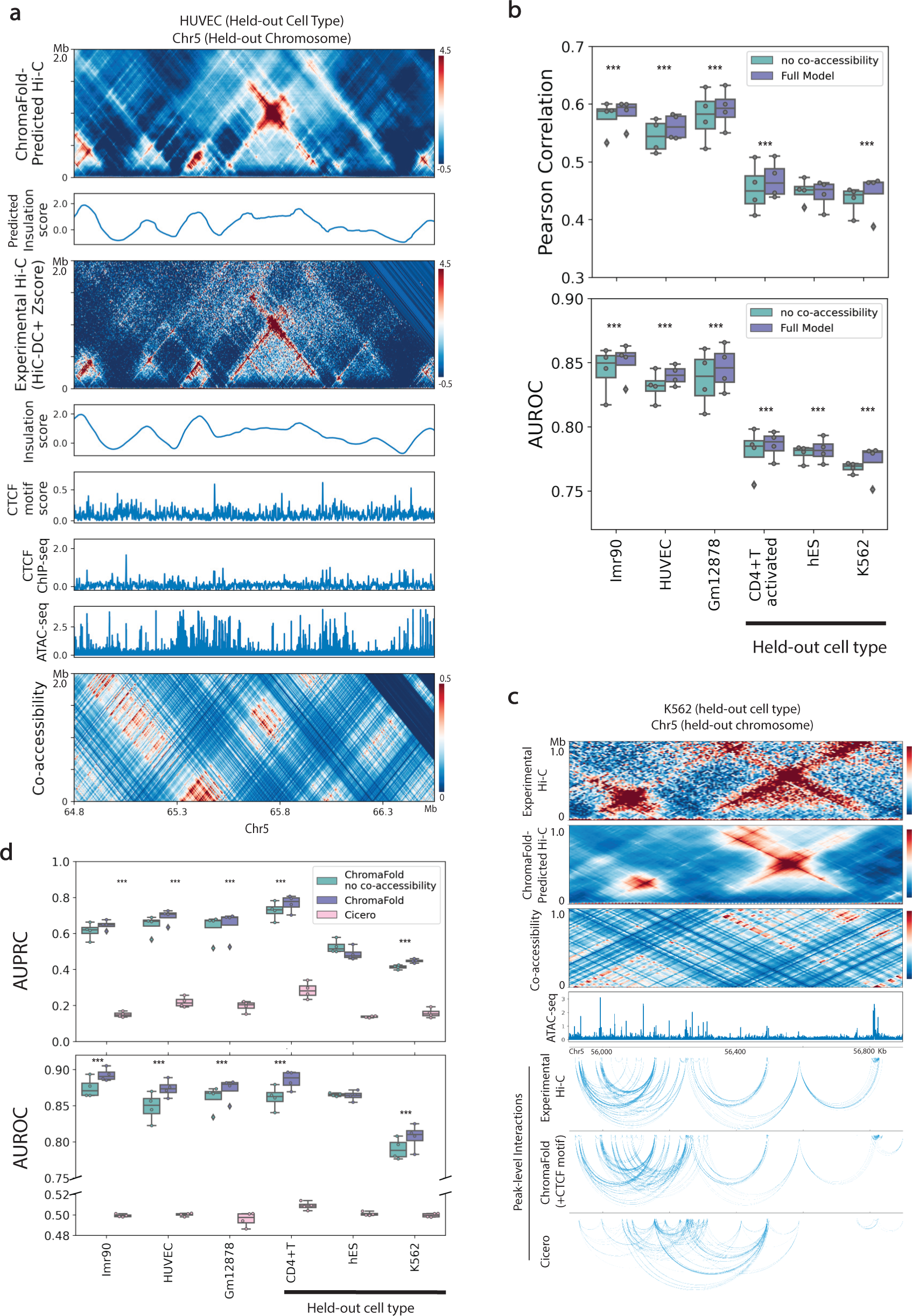
Co-accessibility information improves contact map prediction in new cell types. **a.** Visualization of real vs. ChromaFold-predicted Hi-C contact map, insulation scores, epigenetic tracks, and co-accessibility on held-out chromosome 5 in HUVEC. **b.** Quantitative evaluation of Hi-C map prediction performance by ChromaFold, with and without the co-accessibility input, across training and held-out human cell types/tissues. Box plots show (top) the averaged distance-stratified Pearson correlation between the experimental and predicted contact map and (bottom) the averaged distance-stratified AUROC of significant interactions (top 10% in Z-score), per held-out chromosome. Paired t-test is performed on the distance-stratified person correlation across test chromosomes (P-value: *: <0.05, **: < 0.01, ***: < 0.001). **c.** Visualization of ChromaFold-predicted Hi-C contact map and significant peak-level interactions and Cicero-predicted peak-level interactions in held-out cell type K562 on held-out chromosome 5. **d.** Quantitative evaluation of significant peak-level prediction performance by ChromaFold and Cicero. Box plots show the AUPRC (top) and AUROC (bottom) of significant peak-level interaction prediction per held-out chromosome. Statistical test is the same as above. The paired t-test P-value for both ChromaFold models vs. Cicero are < 0.0001.

### Co-accessibility and CTCF information improve contact map and peak-level interaction prediction

A key goal of ChromaFold is to predict chromatin interactions that connect regulatory elements to their target genes. To this end, we examined the interactions between accessible peaks by associating ATAC-seq peaks with the overlapping genomic bin and calling peak-level interactions based on the experimental/predicted bin-level contact map (**Fig. 2c**, **Methods**). On held-out chromosomes, ChromaFold achieves an average area under the precision-recall curve (AUPRC) of 0.65-0.7 and 0.45-0.75 and an average AUROC of 0.87-0.89 and 0.81-0.89 in training and testing cell types, respectively (**Fig. 2d**). It should be noted that the diminished performance in K562 is likely attributable to the inferior quality of the Hi-C contact map used for evaluation.

We also compared ChromaFold against Cicero, an unsupervised model that first introduced the idea of using co-accessibility to infer chromatin interactions between accessible peaks^18^. Cicero identifies co-accessible pairs of genomic regions based on their correlation in accessibility across metacells, then uses a graphical lasso regularization to predict a sparser contact map. While peaks with high Cicero co-accessibility are indeed enriched for chromatin interactions compared to peaks with co-accessibility < 0, the unsupervised nature of Cicero limits the accuracy of the model, resulting in low precision and recall (**Fig. 2c,d**). Spurious interaction calls are frequently made, since pairs of genomic regions can be correlated in accessibility without representing true 3D interactions (**Fig. 2c**). On the other hand, we also observed numerous examples where interacting regions are uncorrelated across metacells, leading to false negative predictions (**Supplementary Fig. 2c,d**). Additionally, Cicero does not take into account the pseudobulk accessibility profile of peaks and relies solely on correlation structures over metacells, which are heavily influenced by the level of variability in the scATAC-seq dataset (**Supplementary Fig. 1d**). Nevertheless, we did observe a significant improvement in both 3D contact map and peak-level interaction prediction when we incorporated co-accessibility as an input to ChromaFold (**Fig. 2b,d**), suggesting that the supervised model can extract useful information from the co-accessibility signal.

We next compared ChromaFold’s performance when using different types of CTCF information. A qualitative examination of the predicted contact maps in hESC revealed that CTCF information—either predicted binding tracks via motif scores or occupancy from ChIP-seq—is crucial for accurate prediction the contact map (**Supplementary Fig. 2a**). A quantitative analysis of the predicted Hi-C maps and peak-level interactions confirmed this observation, as there was a significant decline in performance when ChromaFold operated without any CTCF information across all tested cell types. The most pronounced performance degradation occurred in hESC, which suggests a potential differential mapping between accessibility, CTCF binding, and chromatin interactions in this cell type. As expected, in the majority of cell types examined, ChromaFold performed optimally when it utilized cell-type-specific CTCF ChIP-seq data in the majority of cell types examined. It should be noted, however, that supplying ChromaFold with predicted CTCF motif information alone was sufficient to significantly enhance its accuracy in predicting both the contact map and significant interactions (**Supplementary Fig. 2b,c**).

As a final method comparison, we benchmarked ChromaFold against C.Origami, a recent model that uses bulk ATAC-seq, DNA sequence, and CTCF ChIP-seq as inputs to predict the 3D contact map^16^. To ensure a fair comparison, we re-trained ChromaFold and C.Origami on the same cell type, IMR-90, with HiC-DC+ Z-score normalized Hi-C contact maps as the target and used the same chromosomes for training, validation (Chr10) and testing (Chr15). While ChromaFold and C.Origami achieved similar performance on the held-out chromosome in the training cell type (**Supplementary Fig. 3a-c**), ChromaFold models outperformed C.Origami on a new cell type, GM12878 (**Fig. 3**). Further expanding our comparison to include two additional cell types used in C.Origami’s cross-cell-type prediction evaluation, K562 and hESC, we found that the ChromaFold model consistently surpassed C.Origami across all metrics when CTCF ChIP-seq data was provided, and achieve comparable performance when using CTCF motif information. Given that HiC-DC+ normalization employs negative binomial regression to control for genomic distance as well as other covariates such as GC content and mappability to identify statistically significant interactions, we propose that this normalization makes contact map prediction more challenging than other normalization methods, such as ICE^33^. Consequently, more heavily parameterized models, like C.Origami, may be more susceptible to overfitting, thereby compromising generalizability.

**Figure 3.**
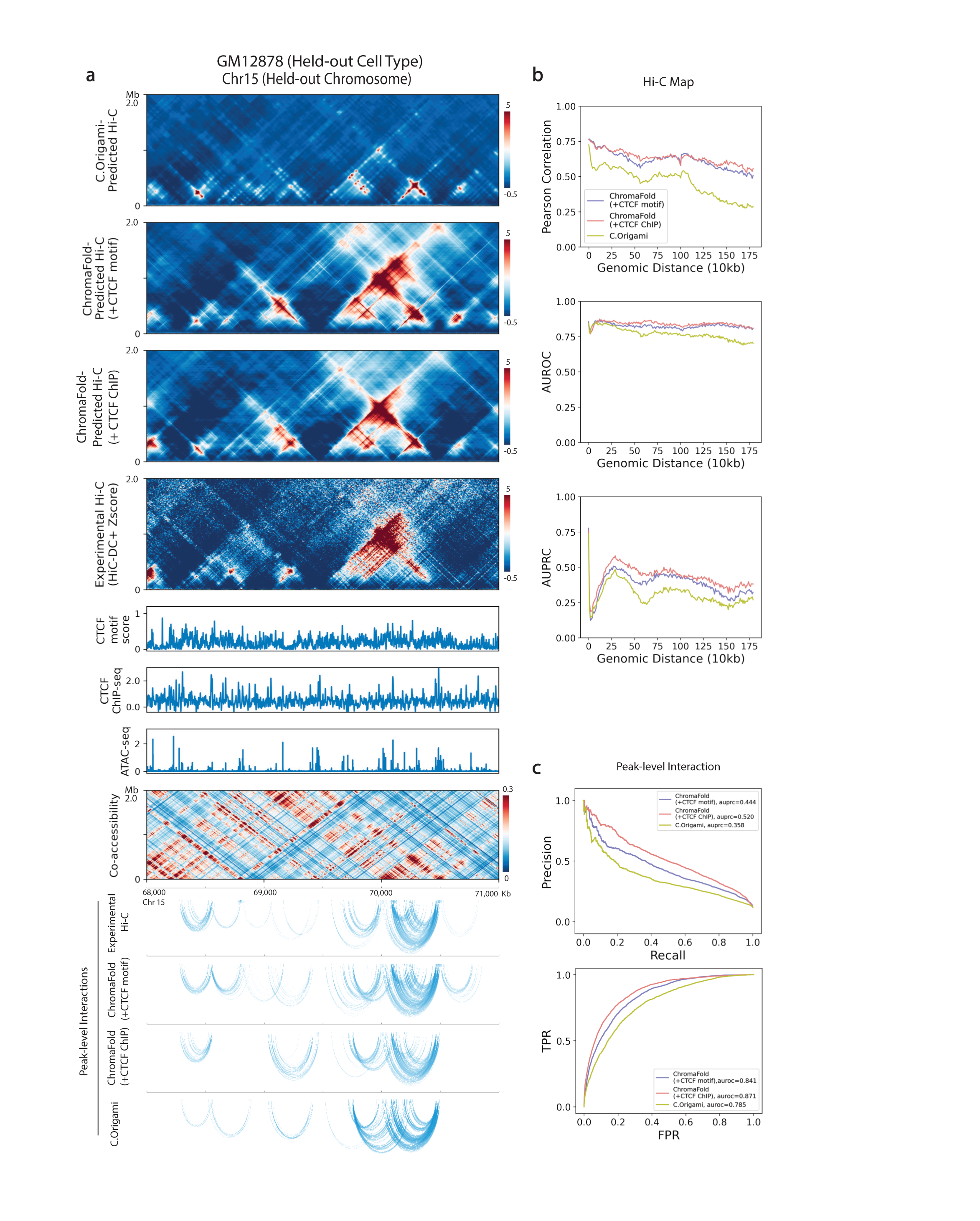
ChromaFold achieves state-of-the-art performance for predicting significant Hi-C interactions in new cell types. C.Origami and ChromaFold were trained using the same training/test chromosomes on IMR-90 to predict contact maps normalized by HiC-DC+ Z-score. **a.** Visualization of C.Origami and ChromaFold-predicted Hi-C contact maps and peak-level interactions in held-out cell type GM12878. **b.** Line plots show distance stratified (top) Pearson correlation between the experimental and predicted contact map, (middle) AUROC and (bottom) AUPRC of significant interactions (top 10% in Z-score) for ChromaFold and C.Origami on held-out chromosome 15. **c.** Line plots show (top) PR curves and (bottom) ROC curves for peak-level interaction prediction on held-out chromosome 15.

### ChromaFold can generalize across species and make cell type-specific predictions

Having shown that ChromaFold can generalize to new human cell types, we proceeded to test whether the model could generalize to a different mammalian genome, since we expect evolutionarily conserved rules governing the mapping between chromatin accessibility and 3D interactions. We therefore directly applied ChromaFold, trained on three human cell types/tissues, to mouse germinal center B cells (GCBs), hematopoietic stem cells (HSCs) and regulatory T (Treg) cells, and evaluated both the predicted contact maps and peak-level interactions. We observed performance comparable to that in human cell types, despite evaluating in a different genome and against lower quality ground-truth Hi-C contact maps in mouse cell types (**Fig. 4c,d**, **Supplementary Fig. 4a,b**). Similar to observations in human test cell types, ChromaFold predictions in mouse are compromised when we ablate the co-accessibility or CTCF motif score input (**Supplementary Fig. 4a**). Notably, we achieve good performance on GCBs with only ∼1500 cells in the scATAC-seq dataset, whereas the smallest training cell type contains ∼3300 cells. These findings suggest that ChromaFold, trained on human data, can generalize to mouse and potentially to other mammalian genomes.

**Figure 4.**
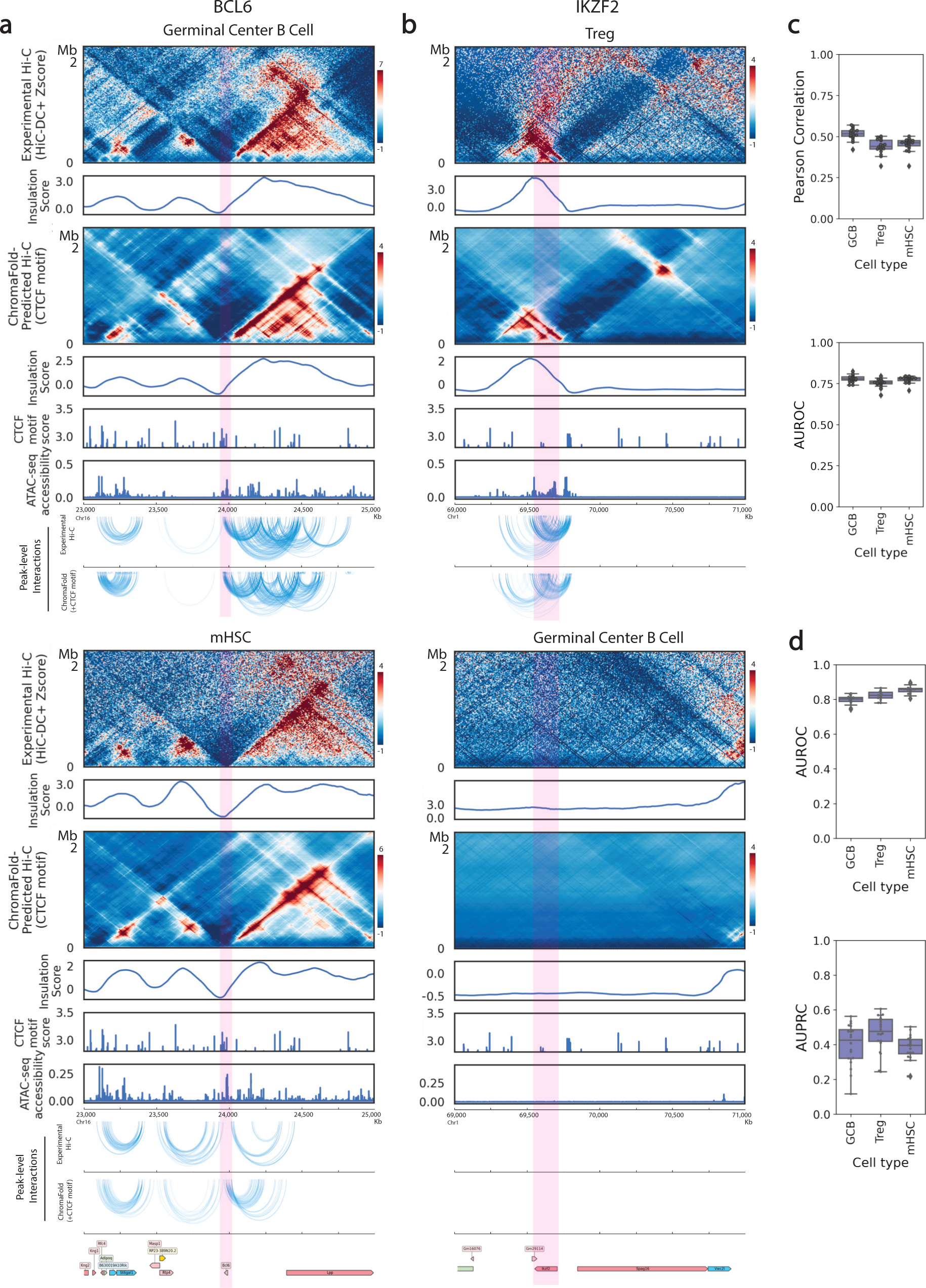
ChromaFold accurately generalizes across cell types and species. **a, b.** Comparison of experimental vs. ChromaFold-predicted Hi-C contact map and peak-level interactions at different loci in the mouse genome across different murine cell types: the *Bcl6* gene locus in mouse germinal center B cells (**a**, top) and in mHSC (**a**, bottom) and the *Ikzf2* gene locus in regulatory T cells (**b**, top) and germinal center B cells (**b**, bottom). **c.** Box plots show (top) the averaged distance-stratified Pearson correlation and AUROC of significant interactions (bottom; top 10% in Z-score), per held-out chromosome across mouse cell types. **d.** Box plots show the AUPRC (top) and AUROC (bottom) of significant peak-level interaction prediction per held-out chromosome across mouse cell types.

Next, we sought to confirm ChromaFold’s ability to make cell-type-specific predictions at loci of interest. Although the predicted CTCF motif score is not cell-type-specific, we expected that the accessibility inputs would confer cell-type-specificity. To illustrate this, we zoomed in on two genes of interest in these cell types: B cell lymphoma 6 (*Bcl6*) and Helios (*Ikzf2*). The *Bcl6* gene encodes a transcription factor that is critical for GCB development^34, 35^. Upon comparing the 3D contact maps at the *Bcl6* locus in GCBs and in HSCs, we observed various conformation changes upstream of the *Bcl6* gene. These differences were accurately captured by ChromaFold-predicted contact maps and insulation scores (**Fig. 4a**). The *Ikzf2* gene is a transcription factor that is essential for the development and function of thymically-derived Treg cells^36, 37^. ChromaFold can predict the presence of chromatin interactions or lack thereof near the *Ikzf2* locus in Treg cells and GCBs, respectively (**Fig. 4b**). Taken together, we conclude that ChromaFold is able to leverage cell-type-specific single-cell chromatin accessibility data and make cell-type-specific contact map predictions.

### ChromaFold can deconvolve chromatin interactions in complex tissue

The ability to study chromatin interactions in fine-grained cell populations can help dissect cell-type-specific gene regulatory programs and contribute to elucidating the pathogenesis of complex genetic diseases. However, the application of experimental techniques such as Hi-C is challenging in rare cell populations due to the difficulty of acquiring sufficient cells for the assay. Although single-cell Hi-C sequencing has made significant advances, the associated experiments remain difficult and expensive, and the sparse contact maps produced are typically analyzed at coarse resolution (100kb-1Mb bins)^11, 38^.

We therefore sought to use ChromaFold to deconvolve chromatin interactions in complex tissues. In scenarios where we possess scATAC-seq and a bulk Hi-C contact map of a tissue or cell population with diverse cell types/states, we decided to fine-tune the pretrained ChromaFold model using input and output data from the mixed population to adapt to the data set. We then applied the fine-tuned model to individual cell populations (clusters) to predict cluster-specific contact maps and thus achieve bulk Hi-C deconvolution.

To evaluate this approach, we applied ChromaFold to deconvolve chromatin interactions in alpha and beta cells within pancreatic islet cell populations using scATAC and bulk Hi-C from non-diabetic islet donors^21^. The predictions were validated against an independent data set containing Hi-C in sorted alpha and beta cells^39^. Our results show that ChromaFold can accurately deconvolve chromatin interactions in the held-out chromosomes (**Fig. 5**). Further, we visualized the predicted interactions at alpha and beta cell marker genes glucagon (*GCG*) and insulin (*INS*). Notably, we predicted a large number of contacts between the *GCG* gene and distal chromatin regions in the alpha cells but not the beta cells, consistent with ground truth data in sorted populations (**Fig. 5a**). On the other hand, we predicted an increased number of contacts between the *INS* gene and both the upstream and downstream chromatin regions in beta cells compared to alpha cells, again matching ground truth contact maps (**Fig. 5b**).

**Figure 5.**
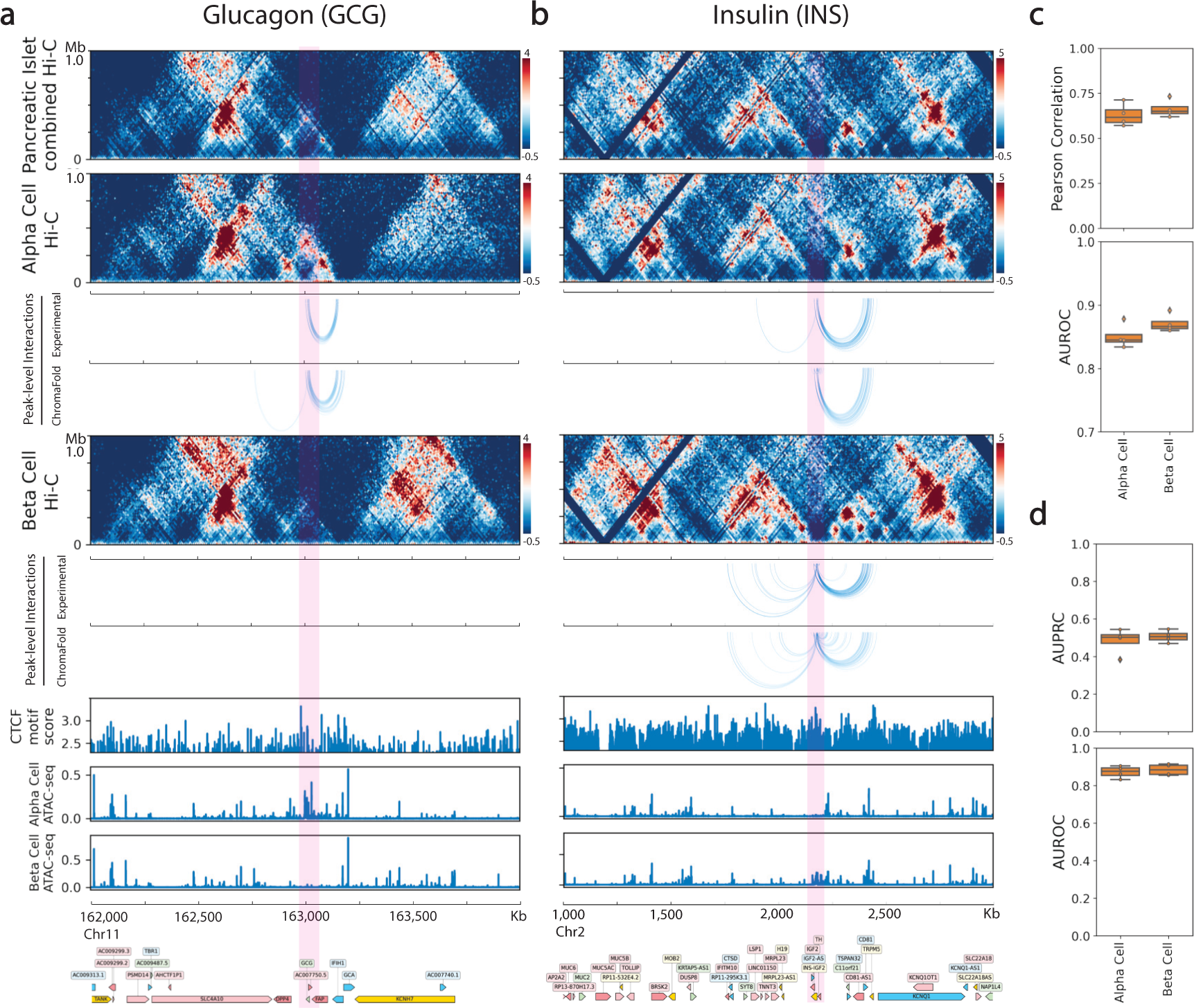
ChromaFold enables deconvolution of Hi-C interactions in pancreatic islet cells. **a, b.** Visualization of peak-level interactions derived from experimental Hi-C data and ChromaFold-predicted Hi-C map in alpha cells and beta cells near the TSS of (**a**) glucagon (*GCG*) and (**b**) insulin (*INS*). **c.** Box plots show (top) the averaged distance-stratified Pearson correlation and AUROC of significant interactions (top 10% in Z-score), per held-out chromosome in alpha and beta cells. **d.** Box plots show the AUPRC (top) and AUROC (bottom) of significant peak-level interaction prediction per held-out chromosome in alpha and beta cells.

## Discussion

Our study demonstrates the utility and potential of ChromaFold for predicting chromatin contacts and mapping putative regulatory elements to their target genes. ChromaFold’s performance, as validated across several metrics and cell types, surpasses previous models such as Cicero and C.Origami, confirming its robustness and versatility. We also found that ChromaFold accurately generalized across species by making cell-type-specific predictions at important loci in diverse mouse cell types from scATAC-seq alone. These findings underscore the shared rules governing the mapping from chromatin accessibility to 3D interaction in mammalian genomes. Furthermore, the ability of ChromaFold to operate on scATAC-seq data sets with ∼1000 cells and the application of ChromaFold for deconvolving bulk contact maps in complex tissues enables the study of chromatin interactions in fine-grained cell populations, providing a novel window into cell-type-specific gene regulatory programs and the dysregulation of these programs in complex genetic diseases.

Our analyses point to several still-unresolved questions for prediction of the 3D contact map: what epigenomic data is most useful for achieving good generalization in new cell types, and what information is captured by DNA sequence models beyond CTCF motif information? Ablation experiments with ChromaFold demonstrated that co-accessibility from scATAC-seq gave a significant performance improvement over pseudobulk accessibility alone. While a number of models including EPCOT^40^ and C.Origami have relied on bulk ATAC-seq as an input signal to help generalization across cell types, our results suggest that covariation in scATAC-seq provides additional information that can be leveraged for contact map prediction. ChromaFold prediction accuracy improved when cell-type-specific CTCF ChIP-seq data was provided as an input.

However, using predicted CTCF motif tracks in place of CTCF ChIP-seq data performed comparably to C.Origami, a state-of-the-art model that uses both a full DNA sequence model as well as ATAC-seq and CTCF ChIP-seq. This result suggests that an improved method for predicting cell-type-specific CTCF ChIP-seq occupancy—in place of the fixed CTCF motif tracks currently used as input—could increase ChromaFold’s accuracy. However, it remains unclear what biological information is captured by introducing a full deep sequence model for contact map prediction, or whether overfitting to spurious sequence signals may be masking relevant information beyond CTCF-associated binding motifs. These questions may be addressed in the coming years through advances in deep learning model interpretation and through ongoing modeling efforts in regulatory genomics. For now, ChromaFold provides a highly favorable trade-off between model complexity, performance, and ease of use, through a lightweight deep learning model that achieves state-of-the-art chromatin map and regulatory interaction prediction from scATAC-seq alone.

## Code availability

Source code is available at: https://github.com/viannegao/ChromaFold/tree/main

## Data availability

Hi-C and scATAC-seq data will be available in GEO.

## Acknowledgements

This work was supported in part by NIH U01 awards HG012103 and DK128852.

M.A.R is supported by a Junior Faculty Scholar Award from the American Society of Hematology. H.L is supported by NYSTEM training award contract C32599GG, and K99 DK128602-01. W.B. is an American Society of Hematology Junior Faculty Scholar Awardee, supported by NIH R01CA270245 and grants from The Leukemia & Lymphoma Society, Lymphoma Research Foundation, and The Follicular Lymphoma Foundation.

## Methods

### Pre-processing of Hi-C and Micro-C data

We used 9 human and 3 mouse datasets (**Supp. Table 1**). For datasets provided in this study and those where a processed .hic file is not available online, Hi-C FASTQ files were aligned to hg38, hg19 or mm10 genomes and reads that are duplicates or invalid ligation products were filtered out using the HiC-Pro^41^ pipeline (v3.1.0) with default settings. Hi-C contact matrices were binned at 10 kb resolution and normalized using the following approaches. ICE normalized contact maps were calculated using the HiCExplorer^42^ package. The counts were log2 normalized using a pseudo-count of 1. Z-score normalization was calculated by the HiC-DC+^26^ package. Specifically, HiC-DC+ models observed raw counts for interaction bins using negative binomial regression to estimate the expected count based on genomic distance, GC content, mappability, and effective bin size based on RE sites in the corresponding pair of genomic intervals.

### Pre-processing of scATAC-seq data

For datasets provided in this study and those where the processed scATAC-seq fragment file was not available online, scATAC-seq FASTQ files were aligned to hg38, hg19 or mm10 and counted by Cell Ranger ATAC v1.2.0^43^ with default parameters. Arrow files were created from the scATAC-seq fragments using ArchR v1.0.1^44^. Specifically, we binarized sparse accessibility matrices binned into 500bp tiles across the genome. Cells with fewer than 1,000 fragments and TSS < 4 were filtered out. Latent Semantic Indexing (LSI) was performed on the 25,000 top variable tiles identified after two iterations of ‘IterativeLSI’ by ArchR. Tiles from non-standard chromosomes, chrM, and chrY were not included. Cells were clustered (method=Seurat, k.param = 30, resolution = 1) and visualized with UMAP^45^ (nNeighbors = 30) using 30 LSI components. For datasets with multiple cell types, we annotated and extracted the cell type of interest by computing the mean gene score of marker genes per cluster. This was cross-checked with cell type annotations provided by the original sources, if available.

### Peak calling

For peak calling of the scATAC-seq data, filtered fragments for cells in each dataset/cell population were aggregated and used as input to the MACS2^46^ peak caller (parameters-f BED, -g 2.7e9, -no-model, -shift −75, -extsize 150, -q 0.05). Peaks were filtered using an IDR^47^ cutoff of 0.05. Peaks within 500bp of each other were merged. A peak-by-cell count matrix was then created by ArchR.

### Bulk ATAC-seq data processing

Bulk ATAC-seq data were obtained from ENCODE^48^ in the form of bam files. Bam files from replicates were merged using samtools^49^, binned at 1bp resolution for C.Origami, and RPKM normalized using the bamCoverage function in deepTools^50^ to generate bigwig files.

### CTCF ChIP-seq and motif score data processing

We obtained the CTCF motif scores from the *CTCF* R package^27^, an AnnotationHub resource that represents genomic coordinates of FIMO-predicted CTCF binding sites for human and mouse genomes. Specifically, CTCF motif scores were generated by scanning for all three JASPAR^28^ CTCF PWMs in genomic DNA sequence using FIMO^25^. CTCF ChIP-seq data were obtained from ENCODE in the form of bam files. Bam files from replicates were merged using samtools, binned at 50bp resolution for ChromaFold and 1bp resolution for C.Origami, and RPKM normalized using the bamCoverage function in deepTools to generate bigwig files. The log2 fold change from the control ChIP-seq in the corresponding cell types were computed using the bigwigCompare function in deepTools.

### ChromaFold input data processing

ChromaFold takes three inputs: pseudobulk chromatin accessibility, co-accessibility profiles across cells, and predicted CTCF motif score/CTCF ChIP-seq. The pseudobulk chromatin accessibility is obtained by aggregating the accessibility profile across single cells in a population binned at 50bp, library-size normalizing and log transforming with a pseudocount of 1. The co-accessibility is derived by first generating metacells to combat sparsity in scATAC-seq datasets, then calculating the Jaccard similarity between binarized accessibility profiles across metacells, binned at 500bp. Metacells are generated using the same algorithm used by Cicero^18^. The CTCF motif score for each 50bp bin in the genome is defined as the maximum score assigned to any genomic region that overlaps at least 10bp with the 50bp bin.

### ChromaFold model architecture

The ChromaFold model consists of two feature extractors and a linear predictor module. The first feature extractor takes the pseudobulk accessibility and the CTCF motif score or ChIP-seq signal as two channels. This feature extractor consists of fifteen 1D convolutional layers followed by batch normalization and ReLU activation. Next, we perform outer-concatenation where the model transforms the resulting L x C matrix, where L is the length of the output vector and C is the number of channels, into a L x L x 2C by performing point-wise concatenation of the output features. This operation allows the information from pairs of genomic bins to be joined together. We implement a skip connection with the input layer by average-pooling the input and transforming into a 3D tensor via outer-concatenation. After concatenation, the data is passed through three 2D convolutional layers followed by a linear layer to consolidate the extracted features, producing a latent representation of the two input tracks.

The second feature extractor takes the co-accessibility data as input. For memory efficiency, we only compute the co-accessibility between the bins in the center 10kb region with the rest of the bins in the 4.01Mb region as input. We use three 1D convolutional layers followed by two residual blocks and three additional 1D convolutional layers. Finally, a linear layer consolidates the extracted features and produces a latent representation of the co-accessibility input. These latent representations of the genomic region are concatenated and passed through a final linear layer to predict the contact between genomic bin *t* and its neighboring bins within a 2Mb distance, which corresponds to a V-shaped stripe (V-stripe) in the contact map centered at t.

### ChromaFold model training

We trained ChromaFold using data pooled from three cell types, IMR-90, Gm12878 and HUVEC. Chromosomes 3 and 15 were used for validation, chromosomes 5, 18, 20, 21 were held out for testing and evaluating model performance, and the rest were used for training. For each V-stripe prediction centered at genomic bin *t*, the input is the 4.01Mb region centered at *t*. During training, we randomly subsampled 500-5000 single cells and 400-1000 metacells from the population per iteration for pseudobulk accessibility and co-accessibility computation, respectively. This data-augmentation step was critical for improving model generalizability to datasets of varying quality and size. We injected additional variation into the input by randomly shifting by −50 or 50 bp. Since neither our input nor output data contain directionality information, we further reduce redundancies in our model by predicting only one side of the V-stripe, and we simply reversed the input to predict the other side (shared model weights). To improve model stability, we used a two-step approach and first train ChromaFold’s feature extractor 1 to predict the target contact map by appending a dummy linear predictor at the end. After convergence, we froze the weights for this part of the network while training feature extractor 2 and the final linear module. Genomic regions with low mappability were masked from training based on the total signal for each bin in the contact map. We took the training window to start and end 4 and 5 Mb after the chromosome starting location and before the ending location, respectively, to create buffer regions since ChromaFold requires 4.01Mb windows as inputs. The prediction target is the HiC-DC+ normalized Z-score, with outlier target values clipped to lie between −16 and 16 to avoid training bias. We optimized the MSE loss using stochastic gradient descent. We trained the model for 30 epochs and implemented early stopping with a patience of 10 epochs, learning rate of 1e-6 and weight decay 1e-3. The model was trained on a single NVIDIA Tesla V40 GPU for ∼5 hours when using one training cell type and ∼14 hours when using 3 training cell types.

### De novo contact map prediction in a new cell type

The ChromaFold model trained on IMR-90, Gm12878 and HUVEC can be directly applied to other cell types and species without retraining. To perform de novo contact map prediction, we supplied scATAC-seq data of the new cell type and predicted CTCF motif scores in the corresponding genome to ChromaFold. If CTCF ChIP-seq data was available for the test cell type, we could alternatively use the *ChromaFold +CTCF ChIP-seq* model.

### ChromaFold Hi-C contact map prediction

To generate the complete predicted contact map for each chromosome, we first performed inference and predicted the interaction between each genomic bin *t* and all its neighboring bins within a 2Mb distance, producing a V-stripe. Since the input region is 4.01Mb centered at the bin *t*, we zero-padded the input vectors if they extended beyond the chromosome edges. We combined the predicted V-stripes and averaged the two predictions for each genomic bin. Contact map prediction for one full chromosome took on average ∼1.5 minutes on a standard GPU like NVIDIA Tesla V40.

### Distance-stratified correlation

To evaluate the overall performance of genome-wide chromatin contact map prediction, we computed the distance-stratified correlation between the experimental and predicted contact maps. The rationale for distance-stratification is to remove any remaining genomic distance effect and avoid inflating the correlation. Specifically, we computed the Pearson correlation for all interactions with genomic distance *d* for *d* from 0 to 2Mb, for each chromosome. We then used a paired t-test^51^ to compare the performance between models. In the boxplot visualizations, each point represents the Pearson correlation averaged across genomic distance, per chromosome.

### Topologically associated domain (TAD) annotations

We called TADs at 10 kb resolution using TopDom^52^ (v0.0.2) using w=30 on normalized Hi-C contact maps and predicted contact maps and used the insulation scores to evaluate ChromaFold’s ability to predict TAD structures.

### Significant interactions

We defined significant interactions at the genomic bin level as interactions with the top 10% HiC-DC+ Z-scores per chromosome. For each chromosome and at each genomic distance (incrementing by 10kb), we used AUROC and AUPRC to evaluate how well significant interaction are captured by ChromaFold’s predicted contact map. We used a paired t-test to compare the performance between models. In the boxplot visualizations, each point represents the corresponding metric averaged across genomic distance, per chromosome. To define significant peak-level interactions, we first mapped each peak to the overlapping genomic bin(s) at 10kb resolution. If a peak overlapped two bins, it was assigned to both. Next, we labeled pairs of peaks as significantly interacting if the corresponding HiC-DC+ FDR-corrected p-value is less than 0.25. The distance-stratified AUROC and AUPRC were computed in a similar fashion as described above.

### Benchmarking against Cicero

We used Cicero to calculate co-accessibility for pairs of peaks. The same metacell groupings used for ChromaFold training/inference were used for running Cicero. We then used Cicero to calculate co-accessibility using a window size of 1 Mb and a distance constraint of 500 kb. We evaluated the performance of peak-level significant interaction prediction using Cicero co-accessibility at various cutoffs and compare with that using ChromaFold-predicted contact maps. All evaluations of peak-level significant interactions were distance-constrained to 500kb for comparison with Cicero.

### Benchmarking against C.Origami

To ensure a fair comparison, we re-trained ChromaFold (with CTCF motif score or with CTCF ChIP-seq) and C.Origami on the same cell type, IMR-90, towards HiC-DC+ normalized Hi-C contact maps and used the same chromosomes for training, validation (Chr10) and testing (Chr15) as specified in C.Origami^16^. The training procedure for ChromaFold was the same as described above, and that for C.Origami was the same as described in the original paper. C.Origami converged after training for 45 epochs. After training, we evaluated the performance of both models on the test chromosome in IMR-90, as well as in three held-out cell types GM12878, K562 and hES. For held-out cell types, we used the IMR-90-trained models but used GM12878/K562/hESC inputs to make de novo contact map predictions. For both models, we merged predictions into a chromosome-wide Hi-C contact map and evaluated the following metrics: 1) distance-stratified Pearson correlation, 2) distance-stratified bin-level significant interaction prediction and 3) peak-level significant interaction prediction.

### Deconvolution of chromatin interactions in alpha and beta cells in the pancreatic islet

ChromaFold can be used for deconvoluting chromatin interactions in complex tissues. Using the scATAC-seq and bulk 3D contact map for pancreatic islet cells, we fine-tuned the pretrained ChromaFold model for 1 epoch on the training chromosomes to better adapt the model predictions to the dataset. We then applied the fine-tuned model to alpha and beta cell populations to achieve deconvolution. Specifically, we extracted the alpha and beta cell clusters from the scATAC-seq to use as input to ChromaFold to generate deconvolved contact map predictions. Next, we used the deconvolved contact maps to generate peak-level interaction predictions as described in the section above. We evaluated the deconvolved chromatin interaction predictions using an independent dataset with Hi-C of sorted human alpha and beta cell populations. For peak-level interaction visualization, we restricted to only interactions involving peaks that lie within 10Kb of the TSS of the highlighted genes. The overall contact map prediction quality was evaluated using distance stratified Pearson correlation. Significant bin- and peak-level interaction predictions were evaluated using distance-stratified AUROC and AUPRC.

### Single-cell ATAC sequencing data collection

Human embryonic stem cells were harvested for single-cell multiome analysis with targeted collection of ∼7000 cells. Nuclei were isolated with Demonstrated Protocol Nuclei Isolation for Single Cell Multiome ATAC+Gene Expression Sequencing_RevA. 500K cells underwent lysis in 500μl lysis buffer in ice for 3 mins then were subjected to the standard protocol for wash and counting. Single-cell Multiome libraries were generated with the 10x Genomics Chromium Next GEM Single Cell Multiome ATAC + Gene Expression Kit following the manufacturer’s guidelines. The libraries were sequenced on the NovaSeq 6000 platform following the manufacturer’s guidelines.

To collect scATAC-seq data in mouse hematopoietic stem cells (Lin-Kit+ cells), bone marrow cells were harvested from total n=3 C57BL6 wildtype mice and subjected to red blood cell lysis. Bone marrow cells were then incubated with MACS beads (CD117, Miltenyi Biotec, 130-091-224). Then enriched c-Kit+ cells were collected by running AutoMACS (Miltenyi Biotec) according to the manufacturer’s instructions. The cells were then stained with cocktail: Lineage marker (CD3, CD8, Gr1, B220, CD19 and Ter119)-PE-Cy5, cKit-APC-Cy7 and DAPI. Live Lin-cKit+ cells were sorted on BD Aria machine. Freshly sorted cells were then resuspended in PBS+0.04% BSA at around 300k/250ul followed by scATACseq protocol.

### Hi-C data collection

Isolation of murine regulatory T cells was conducted as previously described^53^. Cell suspension was made from pooled secondary lymphoid organs (spleen; peripheral and mesenteric lymph nodes) of Foxp3-GFP mice^54^ and CD4 T cells were enriched using the Dynabeads Flowcomp Mouse CD4 Kit (ThermoFisher, 11461D) according to manufacturer’s instructions. The resulting cells were stained with antibodies, washed extensively, resuspended in isolation buffer (PBS w/ 2% FBS, 10 mM HEPES buffer, 1% L-glutamine, and 2 mM EDTA) containing 0.01% SYTOX Blue dead cell stain (ThermoFisher, S34857) to facilitate dead cell exclusion, and sorted on a FACSAria (BD) instrument. Treg cells (TCRβ+CD4+Foxp3-GFP+) and naïve conventional CD4 T cells (TCRβ+CD4+Foxp3-GFP−CD44loCD62Lhi) were sorted by gating on the respective populations. Hi-C was performed as previously described^55^. Briefly, sorted T cell populations (approximately 1×105) were cross-linked in 1% formaldehyde for 10 minutes and quenched in 125mM glycine. Cross-linked cells were lysed and chromatin was restriction enzyme digested using the Arima HiC+ kit (Arima Genomics, San Diego, CA) with manufacturer recommended protocol adaptations for low cell input. Digested and reverse crosslinked DNA was eluted in 100uL and fragmented to 350 bps using a Covaris E220 sonicator (Covaris, Woburn, MA). Sheared genomic material was enriched for biotinylated DNA using streptavidin beads followed by library preparation using Arima protocol modifications for Accel-NGS 2S DNA plus library kit (IDT, Coralville, IA). After end repair and ligation, libraries were quantified using the KAPA library quantification kit (Roche, Indianapolis, IN) and PCR amplified for the number of cycles required to generate >4nM per library. Hi-C libraries were sequenced on an Illumina NovoSeq and raw sequencing data in the Fastq format were obtained.

Germinal Center B cell centrocytes and centroblasts cells were sorted from the spleens of mice immunized with SRBCs for 8 days. Briefly, single-cell suspensions were stained with anti-B220, anti-CD95/Fas and anti-GL7. Centrocytes (Live B220+CD95/Fas+GL7+CXCR4–CD86+) and centroblasts (Live B220+CD95/Fas+GL7+CXCR4+CD86–) were stained with anti-B220, anti-CD95/Fas, anti-GL7, anti-CXCR4 and anti-CD86. DAPI was used for the exclusion of dead cells. Cell sorting was performed in a BD Influx cell sorter in the Weill Cornell Medicine Flow Cytometry Core Facility. Flow-sorted CB and CC were fixed in 1% formaldehyde for 10 min. Fixation was quenched by the addition of 0.125 M glycine for 10 min. In situ Hi-C was performed as described (Rao et al. Cell 2014). Nuclei were permeabilized, and DNA was digested overnight with 100 U DpnII (New England BioLabs). The ends of the restriction fragments were labeled using biotin-14-dATP and ligated in a 1-ml final volume. After reversal of cross-links, ligated DNA was purified and sheared to a length of ∼400 bp, at which point ligation junctions were pulled down with streptavidin beads, DNA fragments were repaired and dA-tailed and Illumina adaptors were ligated. The library was produced by 6–10 cycles of PCR amplification. Sequencing (paired-end, 50 bp) was performed in a HiSeq 2500 Illumina sequencer in the Weill Cornell Medicine Epigenomics Core.

---

**Supplementary Figure 1.**
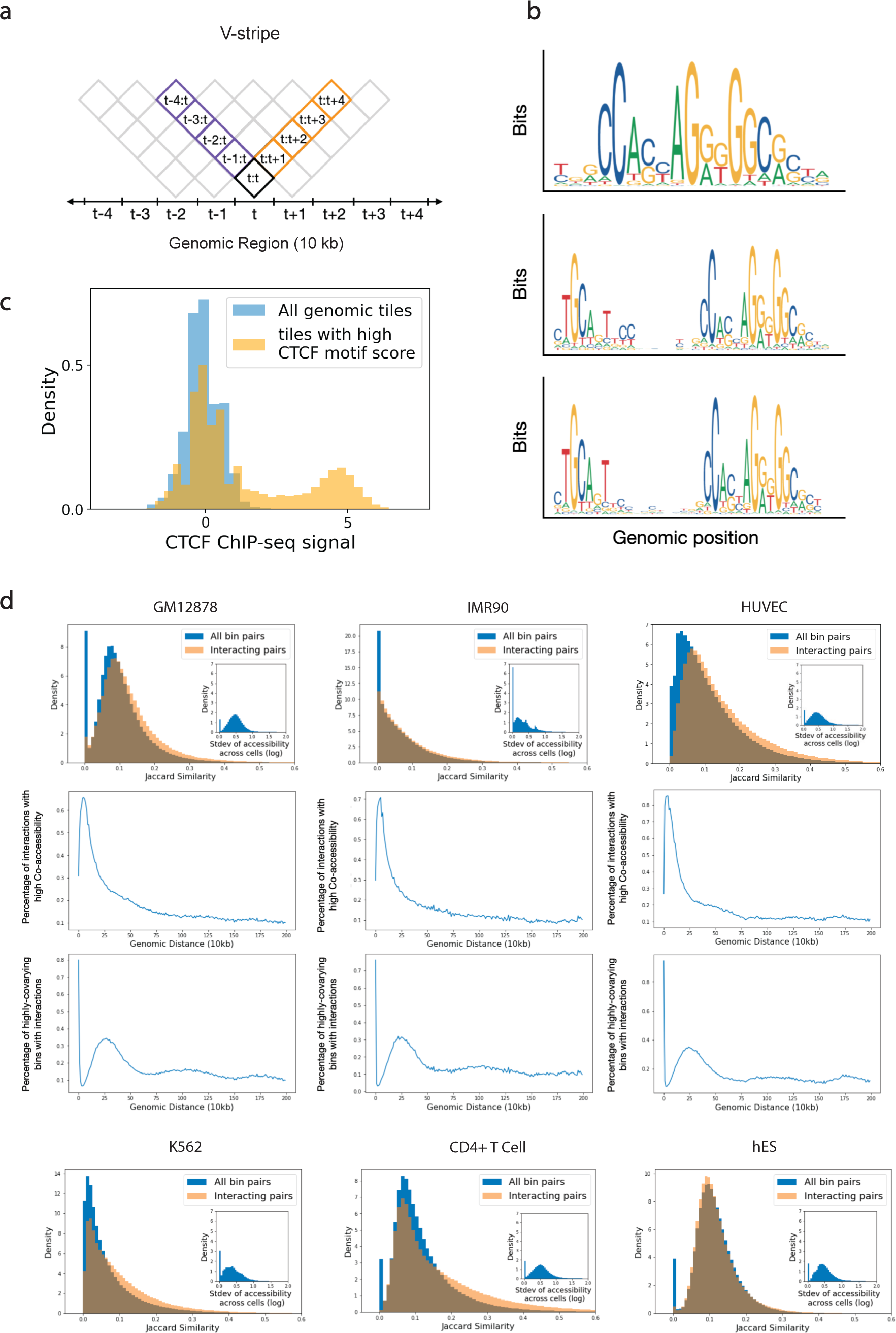
ChromaFold model choices and input analysis. **a.** ChromaFold’s prediction scheme for the chromatin interaction map. Each diamond *t_i*:*t_j* represents the interaction between genomic bins *t_i* and *t_j*. For each input centered around tile *t*, ChromaFold predicts the interaction between tile *t* and its neighboring bins within 2Mb. **b.** CTCF motif PWMs used for CTCF motif scoring. **c.** Histogram shows the distribution of CTCF ChIP-seq signal in genomic bins with top 0.1% CTCF motif score (yellow) and in all genomic bins (blue). **d.** Analysis of the extent of overlap between Hi-C interaction and Jaccard similarity in training cell types. (Top) Histogram show the distribution of Jaccard similarity between interacting bin pairs (top 10% HiC-DC+ Zscore; orange) and all tile pairs within a 2Mb distance (blue). The embedded histogram shows the variability of ATAC-seq accessibility across cells. (Middle) The line plot shows the percentage of interacting bin pairs with high Jaccard similarity (top 10%) at each genomic distance. (Bottom) The line plot shows the percentage of bins with high Jaccard similarity that are interacting at each genomic distance.

**Supplementary Figure 2.**
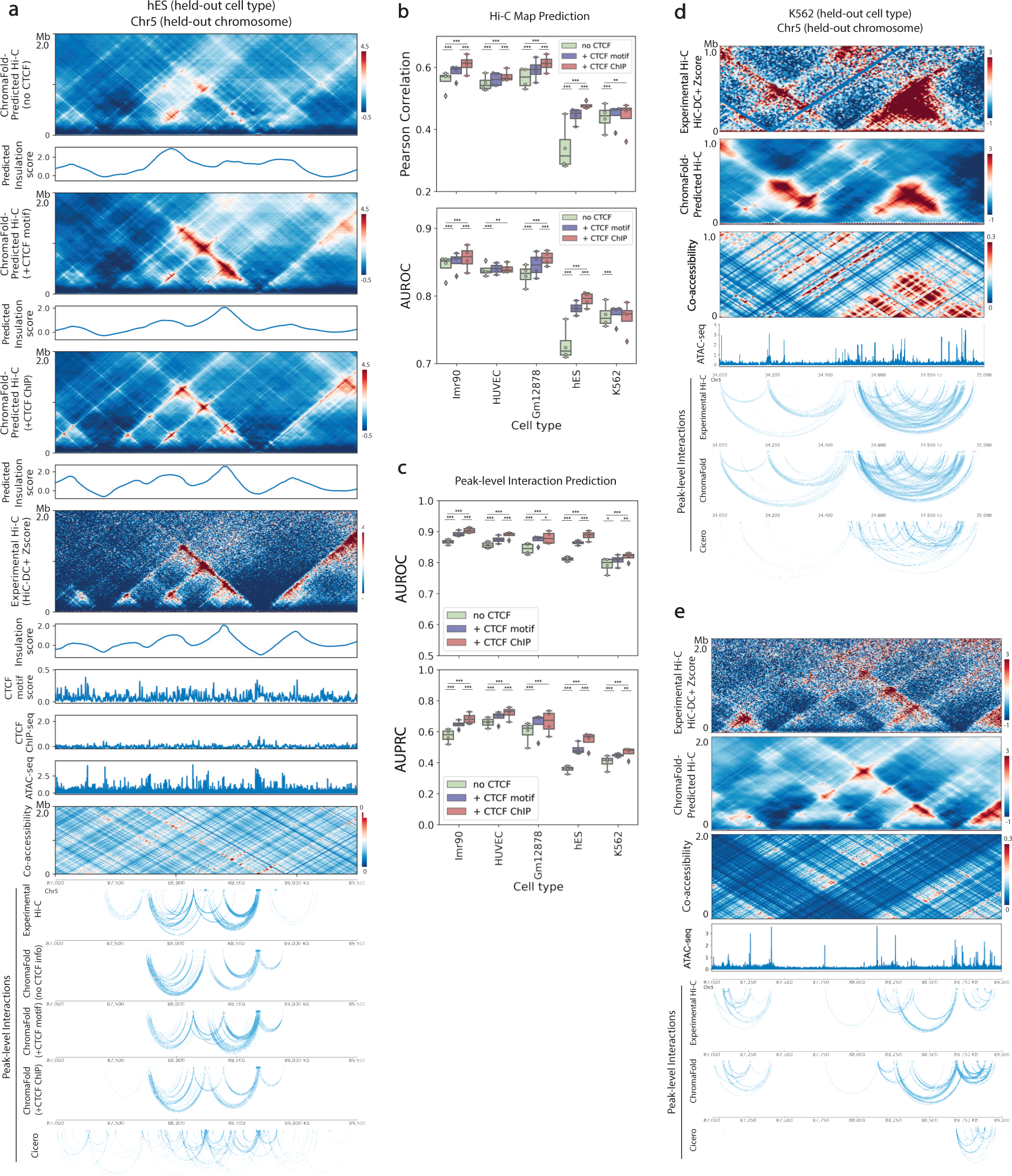
CTCF information is crucial for accurate prediction of Hi-C interactions. **a.** Visualization of Hi-C contact maps, insulation scores and peak-level interactions predicted by ChromaFold using no CTCF information, CTCF motif score and CTCF ChIP-seq data as input in held-out cell type hESC on held-out chromosome 5. **b.** Box plots show (top) the averaged distance-stratified Pearson correlation between the experimental and predicted contact map and (bottom) the averaged distance-stratified AUROC of significant interactions (top 10% in Z-score), per held-out chromosome. Paired t-test is performed on the distance-stratified person correlation across test chromosomes (P-value *: <0.05, **: < 0.01, ***: < 0.001). **c.** Box plots show the AUPRC (top) and AUROC (bottom) of significant peak-level interaction prediction per held-out chromosome. Statistical test is the same as above. **d, e.** Additional visualization of ChromaFold-predicted Hi-C contact map and significant peak-level interactions and Cicero-predicted peak-level interactions in held-out cell type K562 on held-out chromosomes.

**Supplementary Figure 3.**
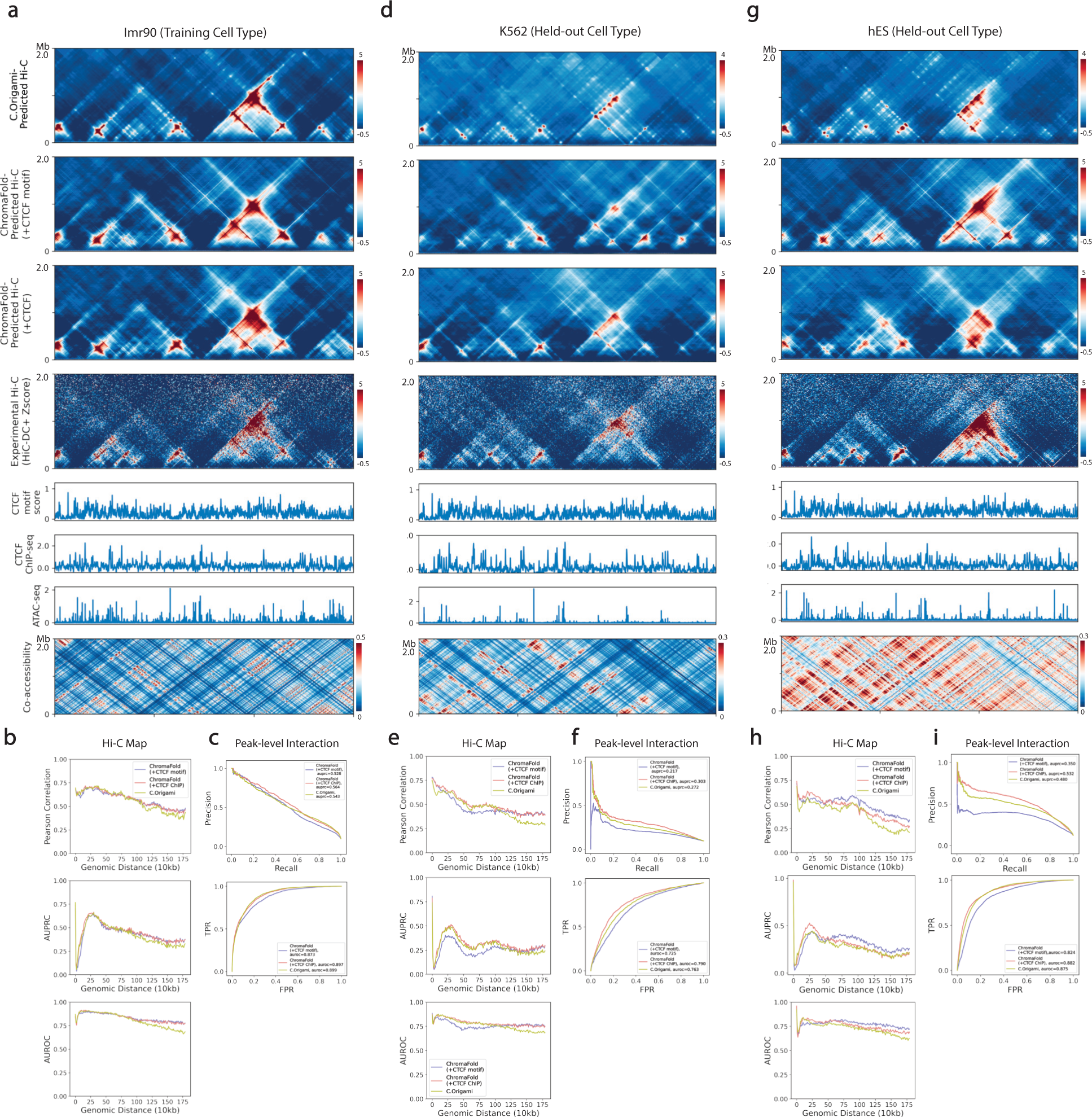
ChromaFold outperforms C.Origami at prediction of HiC-DC+ normalized contact maps in held-out cell types. Visualization of C.Origami and ChromaFold predictions in training cell type IMR-90 (**a, b, c**), held-out cell type K562 (**d, e, f**) and hESC (**g, h, i**): HiC-DC+ normalized Hi-C contact maps (**a, d, g**); distance-stratified Pearson correlation for Hi-C contact map prediction, distance-stratified AUPRC and AUROC of significant interactions (top 10% in Z-score) (**b, e, h**); and PR and ROC curve for peak-level interaction prediction (**c, f, i**).

**Supplementary Figure 4.**
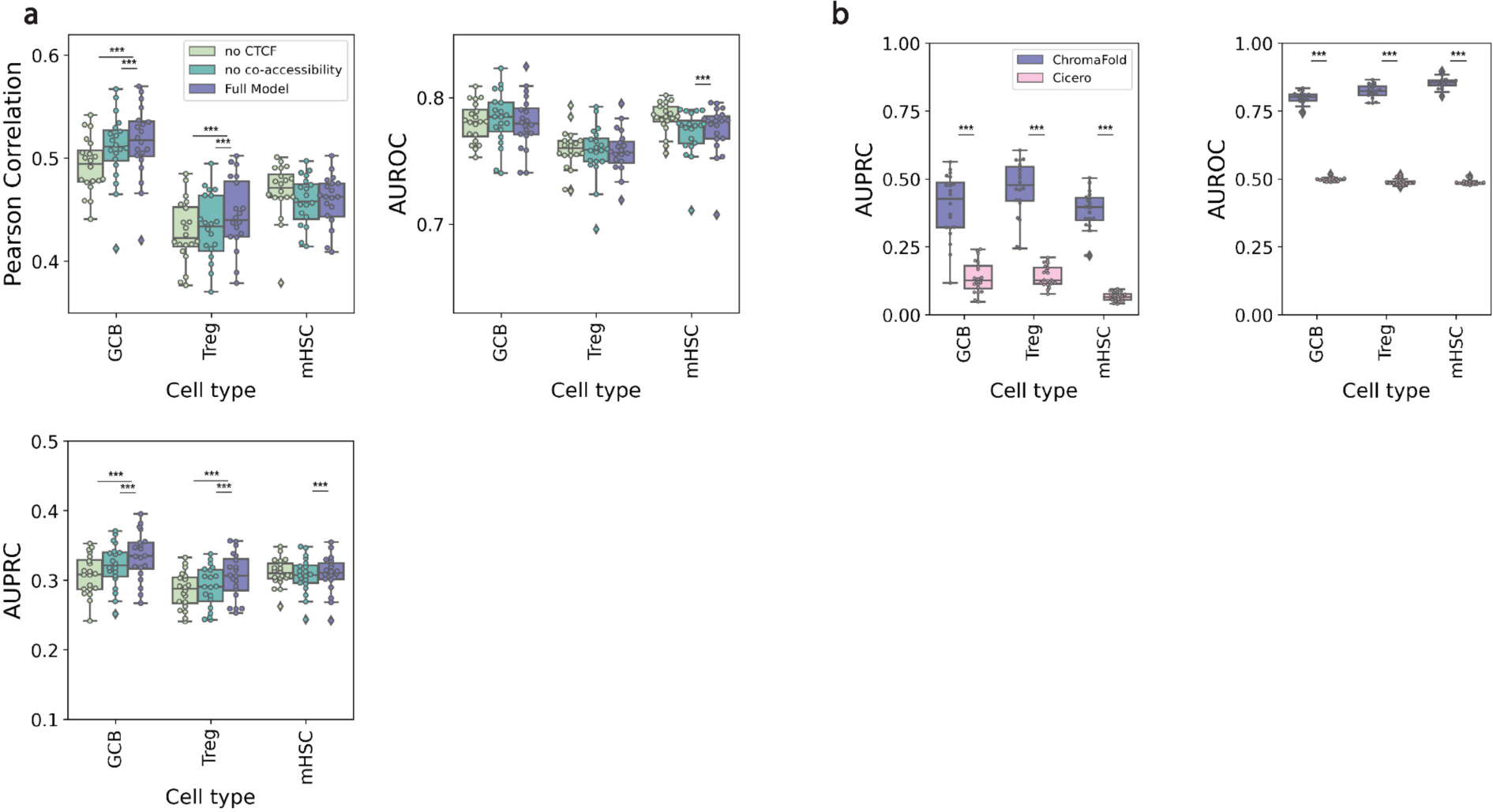
Quantitative evaluation and mode comparison in mouse cell types. **a.** Comparison between ChromaFold model performance in the absence of CTCF motif score information and co-accessibility information. Box plots show the averaged distance-stratified Pearson correlation between the experimental and predicted contact map and the averaged distance-stratified AUPRC and AUROC of significant interactions (top 10% in Z-score), per chromosome. Paired t-test is performed on the distance-stratified person correlation across test chromosomes (P-value *: <0.05, **: < 0.01, ***: < 0.001). **b.** Comparison between ChromaFold and Cicero. Box plots show the AUPRC (top) and AUROC (bottom) of significant peak-level interaction prediction per held-out chromosome. Statistical test is the same as above.

**Supplementary Figure 5.**
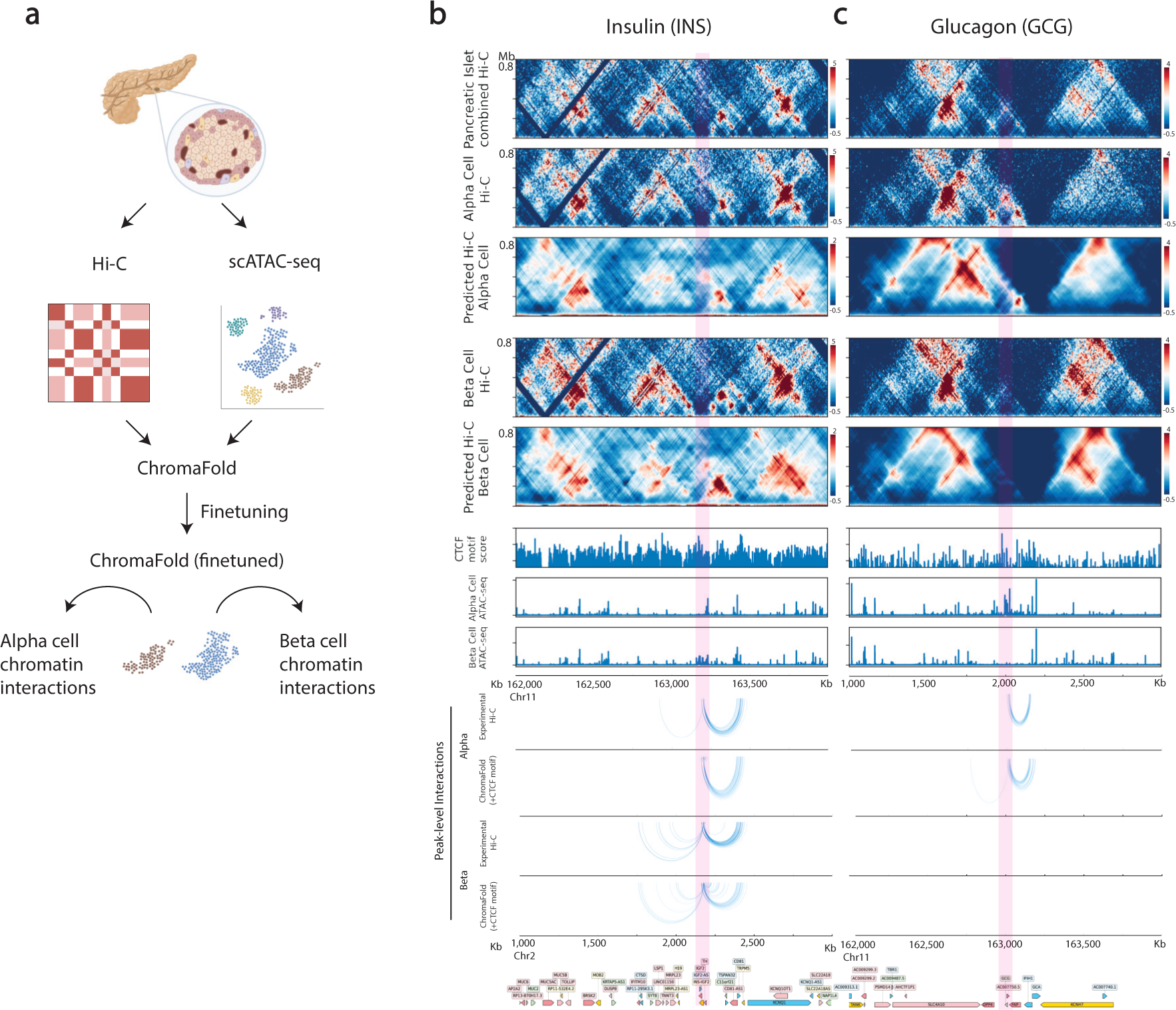
Deconvolution of pancreatic islet cell contact maps at additional loci. **a.** Schematic of ChromaFold applied to the task of deconvoluting chromatin interactions in a complex tissue. **b, c.** Visualization of deconvolved contact maps (top) and peak-level interactions (bottom) in alpha cells and beta cells near the TSS of (**b**) glucagon (*GCG*), and (**c**) insulin (*INS*).

**Table S1:** Summary of data sources for all cell types used in this study.

